# Activation of Cerebellar Lobule VI PC^TH+^–Med Pathway Selectively blocks METH-induced Conditioned Place Preference in mice

**DOI:** 10.1101/2022.08.04.502761

**Authors:** Feifei Ge, Zilin Wang, Guanxiong Wang, Liying Chen, Ping Yang, Jun Wen, Weichao Pan, Wen Yu, Qinglong Cai, Yu Fan, Jiasong Chang, Xiaowei Guan

## Abstract

The cerebellum is implicated in drug addiction. However, the cerebellar neuronal circuitry underlying addiction, especially classic pathway of Purkinje cells (PCs) to deep cerebellar nuclei (DCN), are largely unknown. Here, tracing experiments showed robust projections from the cerebellar lobule VI tyrosine hydroxylase (TH)-positive PCs (PC^TH+^) to CaMKII-positive glutamatergic neurons in medial cerebellar nucleus (Med^CaMKII^), forming PC^TH+^–Med^CaMKII^ pathway. Then, mice were subjected to methylamphetamine (METH)-induced conditioned place preference (CPP), showing that METH exposure excited Med^CaMKII^ and inhibited PC^TH+^. Silencing Med^CaMKII^ by tettox suppressed the formation and expression of METH-induced CPP, but produced serious motor coordination deficits. Chemogenetic activation of lobule VI PC^TH+^–Med pathway during METH CPP training blocked the formation and expression of METH-induced CPP without affecting motor coordination, locomotor activity and sucrose reinforcements in mice. Our findings revealed a novel cerebellar lobule VI PC^TH+^–Med^CaMKII^ pathway and pointed out is critical role in encoding METH-preferred behaviors.

## Introduction

Methamphetamine (METH) is a commonly abused addictive psychostimulant. The reinforcing effects of METH, which are experienced as rewarding, has been believed to drive METH-taking or even addiction. In recent decades, accumulating evidences demonstrate that cerebellum is critical for modulating rewarding circuity (Carta, Chen, Schott, Dorizan, & Khodakhah, 2019; Kostadinov & Hausser, 2022; Wagner, Kim, Savall, Schnitzer, & Luo, 2017), by which may involve in the development of psychostimulants addiction. Further, cerebellum exhibits much sensitive response to METH exposures, as indicated by alterations from gene expressions, neuronal numbers and activities, to structural architecture in METH addicts (Jiang et al., 2021) and METH-exposed animals (Eskandarian Boroujeni et al., 2020; Ferrucci et al., 2007; Hamamura et al., 2010; Thanos et al., 2016). Despite increasing evidence for the involvement of cerebellum in psychostimulants-induced toxicity and behavioral abnormality, however, the cerebellar circuits encoding psychostimulants (e.g. METH)-induced reward process remain largely unknown.

The deep cerebellar nuclei (DCN) are the major source of the cerebellar outputs (Xiao, Bornmann, Hatstatt-Burkle, & Scheiffele, 2018), which send direct or indirect projections to reward-related brain regions such as VTA (Beier et al., 2015), striatum (Hoshi, Tremblay, Feger, Carras, & Strick, 2005) and PFC (Middleton & Strick, 2001), implicating its essential role in the process of rewarding. From medial to lateral, DCN are subdivided into the medial (fastigial in human) cerebellar nucleus (Med), interposed cerebellar nucleus (Int), and lateral cerebellar nuclei (Lat, dentate in human) in rodents. Cocaine administrations increased the neuronal activities in the Med of animals. Remarkably, mice subjected to cocaine-paired odour exhibited higher levels of c-Fos in DCN (Carbo-Gas, Vazquez-Sanroman, Gil-Miravet, et al., 2014), while METH exposure over-activated microglia in DCN of rats (Thanos et al., 2016), suggesting a potential engagement of DCN in psychostimulants addiction. However, few studies have explored the coding role of DCN in the development of addiction.

Anatomically, the innervations on the DCN from cerebellar cortex are merely from cerebellar Purkinje cells (PCs) (Alvina, Walter, Kohn, Ellis-Davies, & Khodakhah, 2008). It is generally believed that PCs send GABAnergic innervations on the DCN neurons, forming an inhibitory neuronal pathway (Hirano, 2018). The cerebellar PCs has been abundantly reported that exhibit sensitive responses to psychostimulants. For example, METH exposure reduced the neural numbers of PCs in rodents (Eskandarian Boroujeni et al., 2020), amphetamine (Sorensen, Johnson, & Freedman, 1982) or cocaine (Jimenez-Rivera, Segarra, Jimenez, & Waterhouse, 2000) exposure reduced the spontaneous discharges of PCs, while cocaine treatment increased BDNF levels but decreased the size and density of dendritic spines in PCs (Vazquez-Sanroman et al., 2015). Recently, tyrosine hydroxylase-positive (TH^+^) cells have been identified in PCs (PC^TH+^) both in rodents (Choi et al., 2012; Lee et al., 2006; T. Locke et al., 2020) and humans (Harrison & Montgomery, 2017). Interestingly, METH administration dose-dependently increased TH levels and neural numbers of the cerebellar PC^TH+^ (Ferrucci et al., 2006; Ferrucci et al., 2007), indicating a potential role of PC^TH+^ in coding addiction. However, the projecting structure of PC^TH+^, especially to DCN, and the role of PC^TH+^ and PC^TH+^-DCN pathway in the process of drugs addiction remains unknown.

In this study, we developed METH-induce conditioned place preference (CPP) model in mice to explore the role of DCN and PC^TH+^–DCN pathway in the acquisition (formation) and expression of METH addiction.

## Results

### Lobule VI PC^TH+^ sends direct projections to Med glutamatergic neurons

To identify the projecting structure of PC^TH+^ to DCN, a retrograde tracing study was performed by unilaterally injecting fluorogold (FG) into the Med to map Med-projecting PCs (Figure 1A, B). As shown in Figure 1C-D, FG-positive retrogradely marked cells throughout a vast majority of cerebellar lobules including the simple lobule (Sim), paramedian lobule (PM), Crus 1 and 2. Among all the cerebellar lobules (Figure 1D-G), the lobule VI contains the highest number of FG and TH double-positive cells, which showed relatively higher percentages of total Med-projecting PCs and TH-positive PCs.

**Figure 1.**
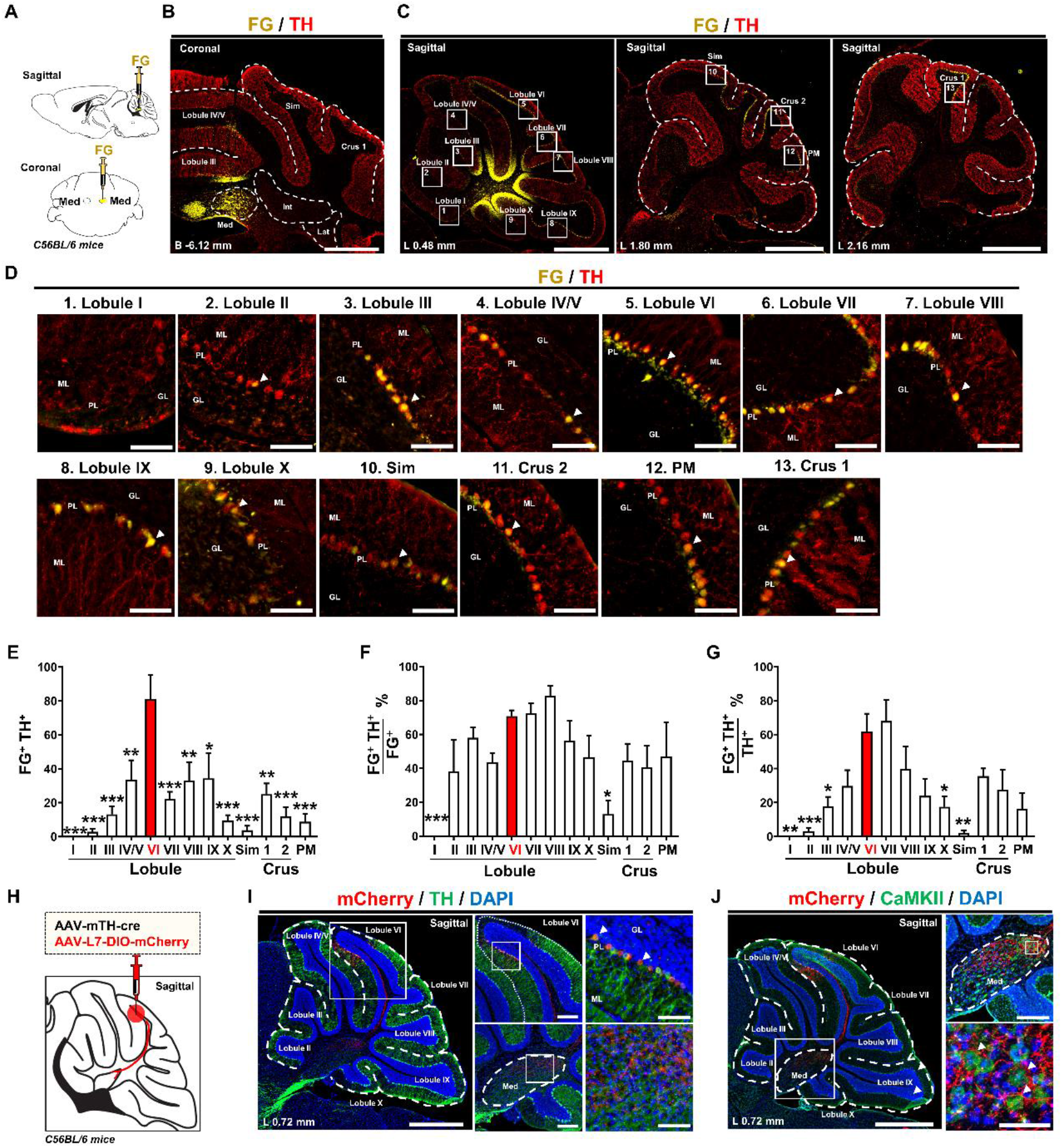
Lobule VI PC^TH+^ sends direct projections to Med glutamatergic neurons. (**A**) Schematics of retrograde tracer FG injection. (**B**) Representative image of FG fluorescence in the Med and immunostaining with TH. Scale bar, 1 mm. (**C, D**) Representative images of the retrograde FG fluorescence in the cerebellar vermis and immunostaining with TH. Scale bar, 1 mm (**C**) and 100 μm (**D**). (**E-G**) Cell counts and the proportions of the FG and TH double positive cells in the cerebellar vermis. One-way ANOVA. F (12,59) = 6.653, *P* < 0.0001. ^*^*P* < 0.05, ^**^*P* < 0.01, ^***^*P* < 0.001 versus lobule VI. n = 4-6. (**H**) Schematics of anterograde tracer virus injection. (**I**) Representative image of mCherry fluorescence in the lobule VI and immunostaining with TH. Scale bar, 1 mm (left), 500 μm (middle) and 300 μm (right). (**J**) Representative images of the anterograde virus-mCherry fluorescence in the Med and immunostaining with CaMKII. Scale bar, 1 mm (left), 500 μm (top right) and 50 μm (bottom right).

Then, an anterograde tracing study were performed by injecting the mCherry-tagged anterograde tracer into the lobule VI to label PC^TH+^ (Figure 1H). As shown in Figure 1I-J, viruses were highly over-lapped with TH staining within lobule VI, and the axon terminals labeled with mCherry from lobule VI PC^TH+^ were abundantly distributed in Med and closely surrounded the CaMKII-positive glutamatergic neurons in Med (Med^CaMKII^).

### METH exposure excites cerebellar Med^CaMKII^ but inhibits lobule VI PC^TH+^

A standard METH-induced CPP procedure were performed in mice (Figure 2A). The c-Fos and NeuN were used as markers to detect the neuronal activities and neurons, respectively. During CPP training, the total distance travelled by mice in CPP apparatus was assessed to reflect the reinforcement effects of METH, by which to some extent can reflect the formation (acquisition) of METH-induced CPP. As shown in Figure 2B, there was no difference in total distance travelled between the METH and saline mice during the Pre-test. During CPP training, METH-exposed mice exhibited extremely higher locomotor activity on METH injection days, but a similar activity on saline injection days (Figure 2C-E), when compared to saline-exposed mice. Then, we observe the expression of METH-induced CPP by examining CPP-test in a subset of mice (Figure supplement 1A). As shown in Figure supplement 1B-D, METH-exposed mice but not saline-exposed mice spend more time in METH-paired chamber than that in Pre-test, and METH injection induced much higher ⊿CPP score than saline injection. As the Med neurons were predominantly glutamatergic (Figure supplement 2), we analyzed the proportions of activated CaMKII-positive glutamatergic neurons (co-expressed with c-Fos) in Med on Day-9 (the last training day of CPP). Compared with the controls, METH-exposed mice showed increased percentage of c-Fos-positive and CaMKII-positive neurons in Med (Figure 2F, G), but similar percentage of that in Lat and Int (Figure supplement 3). However, there was no difference in the c-Fos expression of Med neurons during CPP-test between saline-exposed and METH-exposed mice (Figure supplement 1E, F).

**Figure 2.**
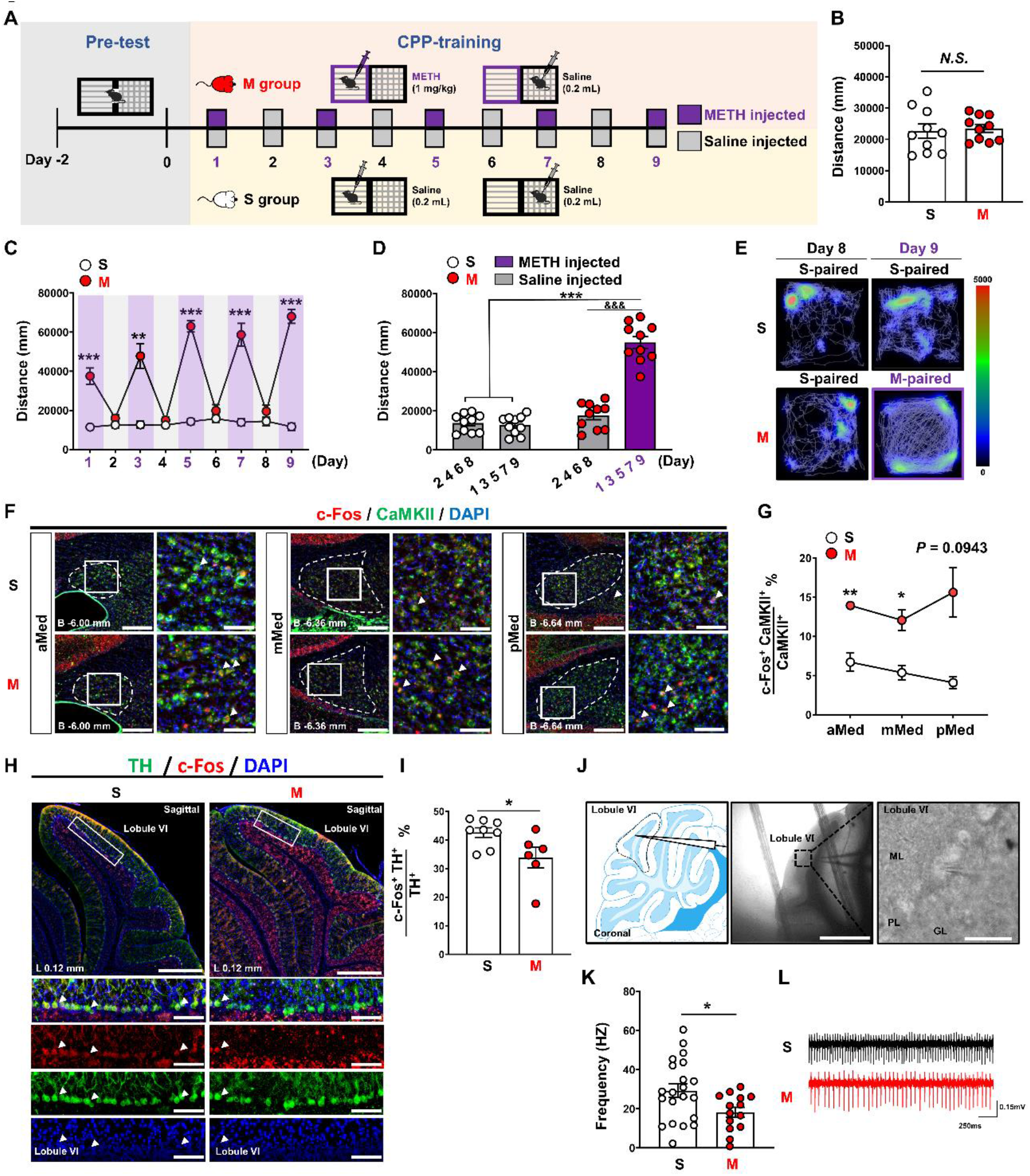
METH exposure excites cerebellar Med^CaMKII^ but inhibits lobule VI PC^TH+^. (**A**) Scheme of METH CPP formation model. (**B**) Distance travelled during the pre-test. Two-tailed unpaired t test, t = 0.3518, df = 18, *P* = 0.7291 versus saline group. n = 10 per group. (**C**) Distance travelled during the CPP formation training. Two-way RM ANOVA. F _treatment × session_ (8,144) = 33.18, *P* < 0.0001. ^**^*P* < 0.01, ^***^*P* < 0.001 versus saline group. n = 10 per group. (**D**) Average distance travelled during METH and Saline injected days. Two-way RM ANOVA. F _treatment × session_ (1,18) = 90.44, *P* < 0.0001. ^***^*P* < 0.001 versus saline group, ^&&&^*P* < 0.001 versus saline injected day of METH group. n = 10 per group. (**E**) Single trial example on the Day 8-9 of METH CPP formation training. (**F**) Immunohistochemistry for c-Fos/CaMKII/DAPI in the Med following METH CPP formation. Scale bar, 300 μm (left) and100 μm (right). (**G**) Percentage of c-Fos and CaMKII double positive cells in the Med^CaMKII^ following METH CPP formation. Two-way RM ANOVA. F _treatment × session_ (2,16) = 1.8, *P* = 0.1958. ^*^*P* < 0.05, ^**^*P* < 0.01 versus saline group. n = 6 per group. (**H**) Immunohistochemistry for c-Fos/TH/DAPI in the lobule VI following METH CPP formation. Scale bar, 1 mm (left) and 100 μm (right). (**I**) Percentage of c-Fos and TH double positive cells in the PC^TH+^ following METH CPP formation. Two-tailed unpaired t test, t = 2.396, df = 12, ^*^*P* = 0.0337 versus saline group. Saline group, n = 8; METH group, n = 6. (**J**) Representative images of electrophysiological recordings on the lobule VI PCs. Scale bar, 1 mm (left) and 50 μm (right). (**K**) Frequency of spontaneous firing in the lobule VI PCs. Two-tailed unpaired t test, t = 2.365, df = 33, ^*^*P* = 0.0241 versus saline group. Saline group, n = 21 cells; METH group, n = 14 cells. (**L**) Sample traces of spontaneous firing in the lobule VI PCs.

Next, the activities of PC^TH+^ in the lobule VI were measured on Day-9. Compared with the saline-exposed mice, METH-exposed mice showed a decrease in c-Fos expression in the lobule VI PC^TH+^ (Figure 2H, J). In parallel, brain slices from METH-exposed mice exhibited a decreased frequency of spontaneous firing in the lobule VI PCs, when compared to that of saline-exposed mice (Figure 2J-L).

### Sustained silencing Med^CaMKII^ before CPP training suppresses METH-induced CPP behaviors and produces motor coordination deficits in mice

To determine the role of Med in METH-induced CPP, virus containing tetanus toxin light chain (tettox) were bilaterally injected into Med to silence Med^CaMKII^ (Zhou et al., 2018) before CPP training (Figure 3A, B). All mice were subjected to METH injection on odd days (Day-1, Day-3, Day-5 and Day-7). As shown in Figure 3C, Viruses were expressed restrictedly and highly over-lapped with CaMKII staining in the Med. As shown in Figure 3D-F of CPP training phase, tettox-injected mice traveled lesser distance in CPP apparatus on METH injection days (Day-1, Day-3, Day-5 and Day-7) when compared with control virus-injected METH mice, and similar distance on saline-injected days to their distance on METH injection days. During CPP test, METH injection increased CPP score in control virus-injected but failed to affect that in tettox-injected mice when compared with the corresponding Pre-test (Figure 3G, H), as well as induced a lower trend of ⊿CPP score in tettox group than control group (Figure 3I). During the Pre-test and CPP-test, there was no difference in the distance travelled between tettox and control group (Figure supplement 4A, B).

**Figure 3.**
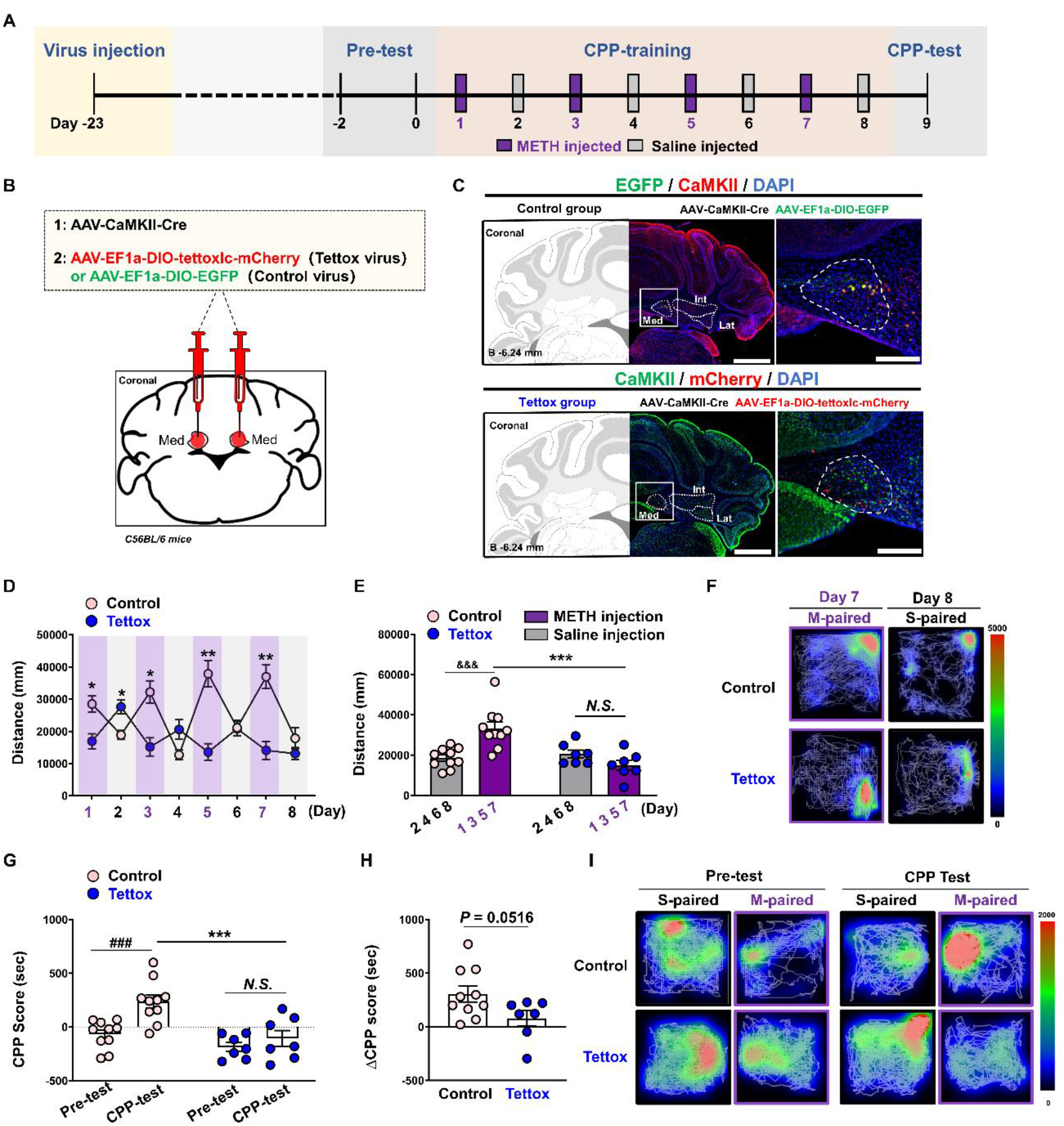
Sustained silencing Med^CaMKII^ before CPP training suppresses METH-induced CPP behaviors. (**A**) Experimental scheme. (**B**) Schematics of virus injection. (**C**) Representative images of fluorescent virus tag (top, EGFP for control group; bottom, mCherry for Tettox group) in the Med and immunostaining with CaMKII and DAPI. Scale bar, 1 mm (left) and 300 μm (right). (**D**) Distance travelled during the CPP formation training. Two-way RM ANOVA. F _treatment × session_ (7,105) = 14.01, *P* < 0.0001. ^*^*P* < 0.05, ^**^*P* < 0.01 versus saline group. Control group, n = 10; Tettox group, n = 7. (**E**) Average distance travelled during METH and Saline injected days. Two-way RM ANOVA. F _treatment × session_ (1,15) =16.05, *P* = 0.0011. ^&&&^*P* < 0.001 versus saline injected day of control group. ^***^*P* < 0.001 versus control group on the METH injected day. Control group, n = 10; Tettox group, n = 7. (**F**) Single trial example on the Day 7-8 of METH CPP formation training. (**G**) CPP scores during the pre-test and CPP test. Two-way RM ANOVA. F _treatment × session_ (1,15) =4.470, *P* = 0.0516. ^###^*P* < 0.001 versus Pre-test of control group. ^***^*P* < 0.01 versus control group during the CPP-test. Control group, n = 10; Tettox group, n = 7. (**H**) ⊿CPP score (CPP test minus pre-test) of the two group. Two-tailed unpaired t test, t = 2.114, df = 15, *P* = 0.0516 versus control group. Control group, n = 10; Tettox group, n = 7. (**I**) Single trial example during the pre-test and CPP test.

A subset of mice was subjected to Rota-rod test, OFT and sucrose-induced SA to examine the effects of tettox in Med^CaMKII^ on motor coordination, general locomotor and natural reward, respectively (Figure 4A). As shown in Figure 4B, tettox-injected mice showed shorter latency to fall than controls. The is no difference in the distance traveled in OFT apparatus between tettox group and controls (Figure 4C, D). In sucrose-trained SA experiments, tettox-injected mice showed occasional increase in active lever presses on Day-10 and Day-11 (Figure 4E) and sucrose water consumption on Day-10 (Figure 4G). However, there was no difference in the lever presses and sucrose consumption during the whole process of sucrose SA between the the two groups (Figure 4E-G).

**Figure 4.**
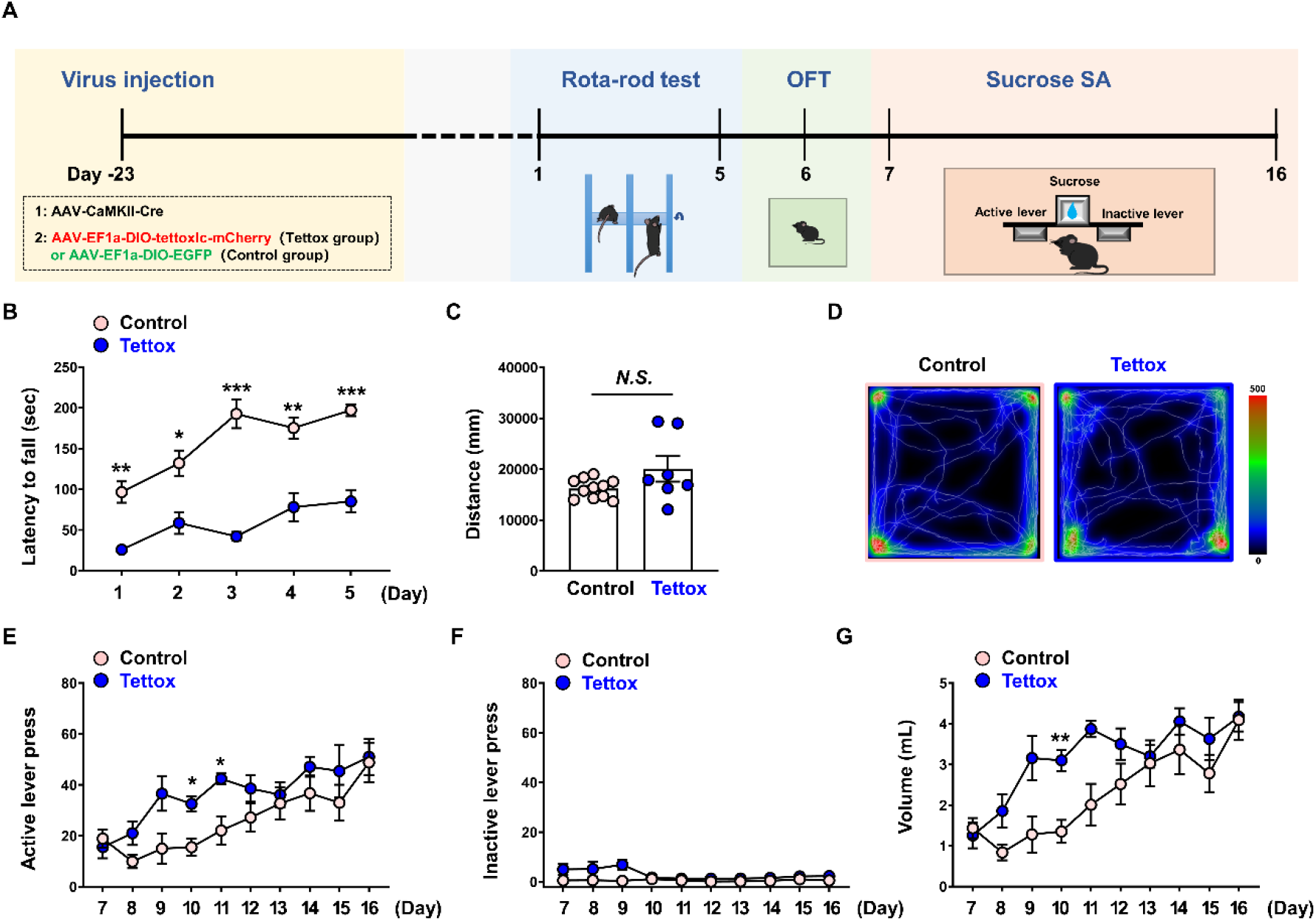
Silencing Med^CaMKII^ produces motor coordination deficits in mice. (**A**) Experimental scheme. (**B**) Latency to fall during the Rota-rod test. Two-way RM ANOVA. F _treatment × session_ (4,64) =5.701, *P* = 0.0005. ^*^*P* < 0.05, ^**^*P* < 0.01, ^***^*P* < 0.001 versus control group. Control group, n = 11; Tettox group, n = 7. (**C**) Distance travelled during the OFT. Two-tailed unpaired t test, t = 1.860, df = 16, _*_*P* = 0.0814 versus control group. Control group, n = 11; Tettox group, n = 7. (**D**) Single trial example during the OFT. (**E**) Number of active lever press during the sucrose reinforcement. Two-way RM ANOVA. F _treatment × session_ (9,144) = 1.220, *P* = 0.2870. ^*^*P* < 0.05 versus control group. Control group, n = 11; Tettox group, n = 7. (**F**) Number of inactive lever press during the sucrose reinforcement. Two-way RM ANOVA. F _treatment × session_ (9,144) = 4.061, *P* = 0.0001. Control group, n = 11; Tettox group, n = 7. (**G**) Volume of sucrose consumption during the sucrose reinforcement. Two-way RM ANOVA. F _treatment × session_ (9,144) = 1.827, *P* = 0.0682. ^**^*P* < 0.01 versus control group. Control group, n = 11; Tettox group, n = 7.

### Chemogenetic activation of lobule VI PC^TH+^–Med pathway during CPP training blocks METH-induced CPP without influencing motor coordination in mice

To examine the role of lobule VI PC^TH+^–Med pathway in METH-induced CPP, AAV encoding chemogenetic activator DREADD (designer receptor exclusively activated by designer drugs; hM3Dq) were bilaterally injected into the lobule VI to modulate PC^TH+^, and the cannulas were placed in the Med. During CPP training, CNO or vehicle were locally infused into the Med 15 min prior to every METH injection day (Day-1, Day-3, Day-5 and Day-7) (Figure 5A, B). As shown in Figure 5C, D, viruses were restrictedly expressed in the lobule VI and highly over-lapped with TH staining, and mCherry-tagged axon terminals from lobule VI PC^TH+^ closely surrounded Med^CaMKII^. As shown in Figure 5E-G of CPP training phase, CNO-treated mice traveled lesser distance in CPP apparatus on METH injection days of Day-5 and Day-7 when compared with vehicle-treated mice, and similar distance on saline-injected days to their distance on METH injection days. During CPP test, METH injection increased CPP score in vehicle-treated but failed to affect that in CNO-treated mice when compared with corresponding Pre-test (Figure 5H, J). The ⊿CPP score in CNO group was much higher than that in vehicle group (Figure 5I).

**Figure 5.**
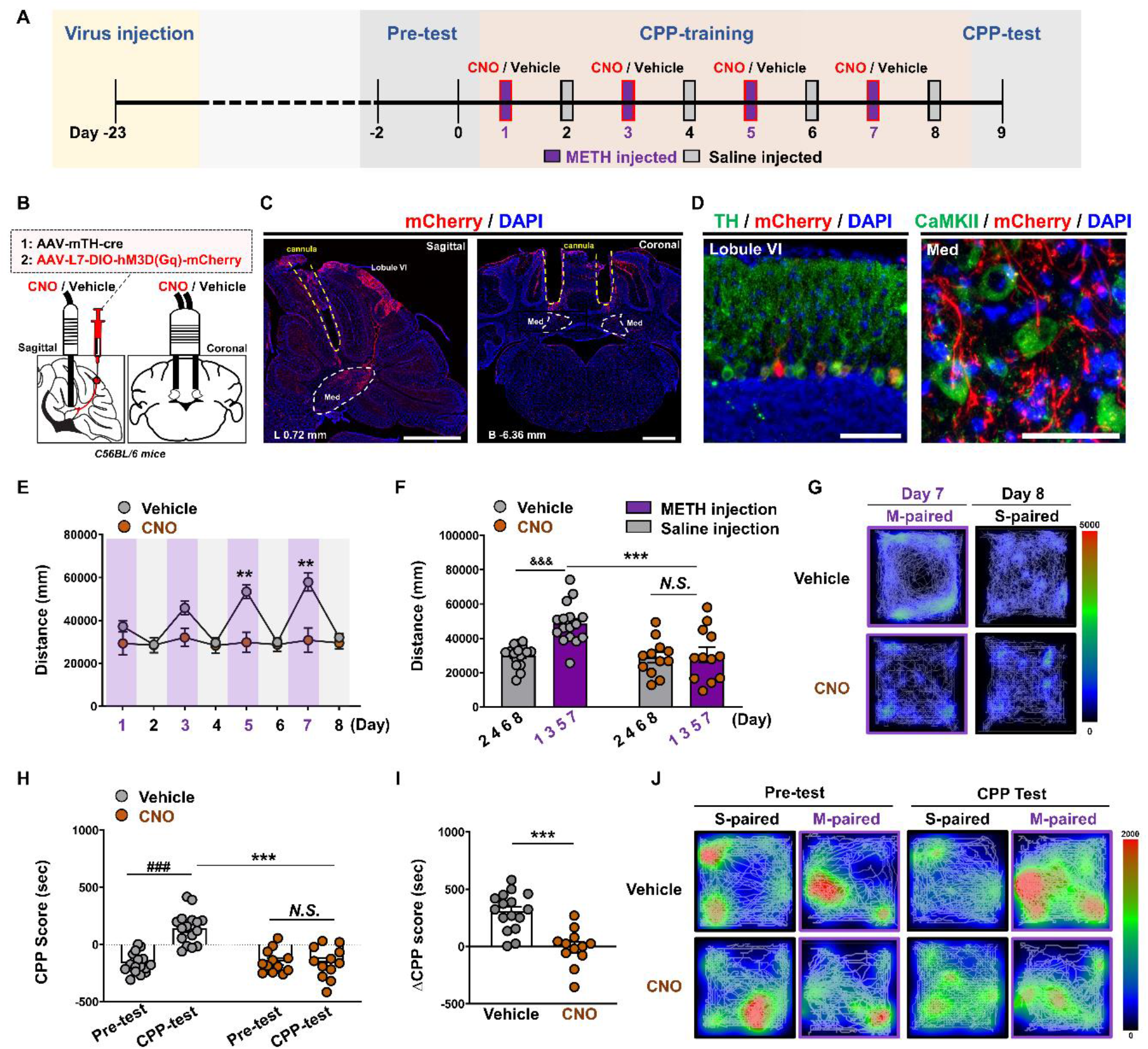
Chemogenetic activation of lobule VI PC^TH+^–Med pathway during training blocks METH-induced CPP in mice. (**A**) Experimental scheme. (**B**) Schematics of virus injection and cannula implantation. (**C**) Representative images of fluorescent virus tag mCherry and cannula implantation in the lobule VI and Med (left, sagittal; right, coronal). Scale bar, 1 mm. (**D**) Representative images of fluorescent virus tag mCherry with TH immunostaining in the lobule VI (left) and with CaMKII immunostaining in the Med (right). Scale bar, 100 μm (left) and 50 μm (right). € Distance travelled during the CPP formation training. Two-way RM ANOVA. F _treatment × session_ (7,182) = 10.88, *P* < 0.0001. ^**^*P* < 0.01 versus saline group. Saline group, n = 16; CNO group, n = 12. (**F**) Average distance travelled during METH and Saline injected days. Two-way RM ANOVA. F _treatment × session_ (1,26) =15.47, *P* = 0.0006. ^&&&^*P* < 0.001 versus saline injected day of saline group. ^***^*P* < 0.001 versus saline group on the METH injected day. Saline group, n = 16; CNO group, n = 12. (**G**) Single trial example on the Day 7-8 of METH CPP formation training. (**H**) CPP scores during the pre-test and CPP test. Two-way RM ANOVA. F _treatment × session_ (1,26) =25.41, *P* < 0.0001. ^###^*P* < 0.001 versus Pre-test of saline group. ^***^*P* < 0.01 versus saine group during the CPP-test. Saline group, n = 16; CNO group, n = 12. (**I**) ⊿CPP score (CPP test minus pre-test) of the two group. Two-tailed unpaired t test, t = 5.041, df = 26, *P* < 0.0001 versus saline group. Saline group, n = 16; CNO group, n = 12. (**J**) Single trial example during the pre-test and CPP test.

A subset of mice was subjected to Rota-rod test, OFT and sucrose-induced SA to examine the effects of lobule VI PC^TH+^–Med pathway in Med on motor coordination, general locomotor and natural reward (Figure 6A). As shown in Figure 6B-D, there was no difference in latency to fall in rod test as well as the distance traveled in OFT apparatus between CNO-treated and vehicle-treated mice. In sucrose-trained SA experiments, there was no difference in active lever presses (Figure 6E), inactive lever presses (Figure 6F), and sucrose consumption (Figure 6G) between the two groups.

**Figure 6.**
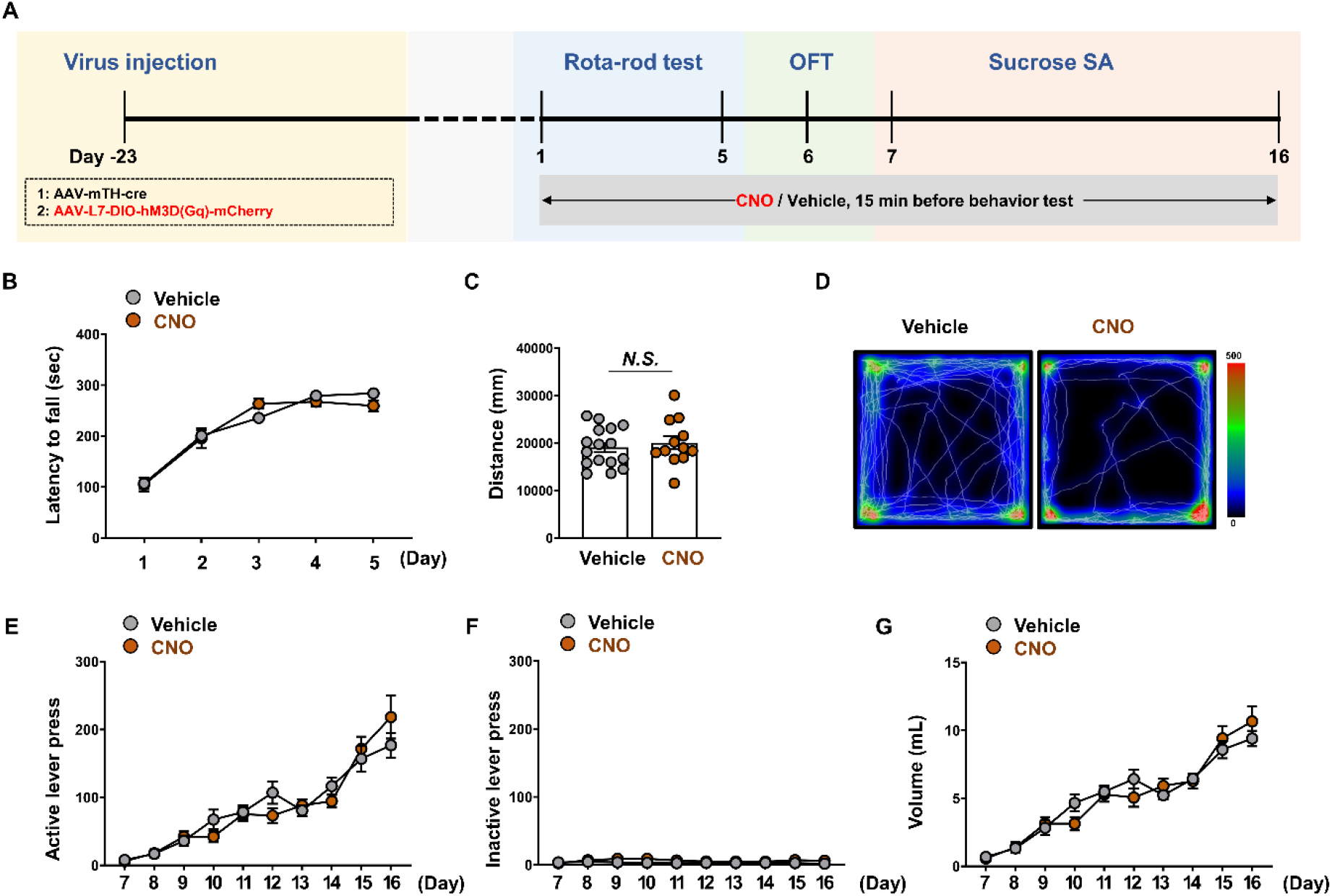
Activation of lobule VI PC^TH+^–Med pathway has no influence on motor coordination in mice. (**A**) Experimental scheme. (**B**) Latency to fall during the Rota-rod test. Two-way RM ANOVA. F _treatment × session_ (4,104) =2.156, *P* = 0.0792. Saline group, n = 16; CNO group, n = 12. (**C**) Distance travelled during the OFT. Two-tailed unpaired t test, t = 0.5942, df = 26, ^*^*P* = 0.5575 versus saline group. Saline group, n = 16; CNO group, n = 12. (**D**) Single trial example during the OFT. (**E**) Number of active lever press during the sucrose reinforcement. Two-way RM ANOVA. F _treatment × session_ (9,234) = 2.276, *P* = 0.0184. Saline group, n = 16; CNO group, n = 12. (**F**) Number of inactive lever press during the sucrose reinforcement. Two-way RM ANOVA. F _treatment × session_ (9,234) = 3.180, *P* = 0.0012. Saline group, n = 16; CNO group, n = 12. (**G**) Volume of sucrose consumption during the sucrose reinforcement. Two-way RM ANOVA. F _treatment × session_ (9,234) = 2.566, *P* = 0.0078. Saline group, n = 16; CNO group, n = 12.

## Discussion

In the last decades, brain regions and their neuronal circuits, such as VTA, nucleus accumbens (NAc), PFC and hippocampus, has been well-studied in the process of addiction (Cooper, Robison, & Mazei-Robison, 2017). Manipulation or even stereotactic destruction of these regions or circuits to some extend alleviate drug-seeking or craving behaviors, but caused strong side-effects on psychiatry abnormalities such as dementia and mental illness. Here, we identified the pathway of cerebellar lobule VI PC^TH+^ to the Med^CaMKII^ might be an idea therapeutic target. Med^CaMKII^ were activated while lobule VI PC^TH+^ were suppressed followed METH CPP formation in mice. Silencing Med^CaMKII^ cells by Tettox during METH CPP training disrupted both the formation and expression of METH CPP, but accompanied by serious deficits in motor coordination. When local activation of the terminals from lobule VI PC^TH+^ within the Med, the formation and expression of METH-induced CPP were totally blocked, importantly with no alteration on the behaviors of natural reward, motor coordination and locomotives.

It is generally believed that drug-related rewarding and drug-associated conditioned memory essentially account for the process of METH addiction. Previous studies showed that the cerebellum is a relevant node for drug-related cue (Gil-Miravet et al., 2021; Jasinska, Stein, Kaiser, Naumer, & Yalachkov, 2014; Li et al., 2015; Moulton, Elman, Becerra, Goldstein, & Borsook, 2014). For instance, cocaine-related stimuli increased activation of the cerebellum in cocaine addicts (Anderson et al., 2006; Grant et al., 1996). Rodent studies showed cocaine-paired olfactory cue increased activity of the granule cell layer of the cerebellar vermis (Carbo-Gas, Vazquez-Sanroman, Aguirre-Manzo, et al., 2014; Carbo-Gas, Vazquez-Sanroman, Gil-Miravet, et al., 2014), and a random cocaine-odour pairing procedure triggered higher cFos expression in the Med (Carbo-Gas, Vazquez-Sanroman, Gil-Miravet, et al., 2014). Non-human primate PCs encoded multiple independent reward-related signals during reinforcement learning (Sendhilnathan, Ipata, & Goldberg, 2021). Here, we found Med^CaMKII^ neurons were notably excited during CPP training, but not CPP test, suggesting Med glutamatergic neurons are responsive to METH instead of METH-paired conditioned stimulus. Importantly, sustained silencing Med^CaMKII^ or activating lobule VI PC^TH+^–Med pathway only during METH CPP training, both blocked METH-induced sensitization behaviors and METH-induced CPP, indicating that Med^CaMKII^ and lobule VI PC^TH+^–Med pathway are involved in coding CPP training process, which are thought of acquisition (or formation) phase of METH-induced CPP.

One recent work has revealed a direct monosynaptic pathway from the cerebellar DCN to the VTA which plays a key role in modulating reward-related behavior (Carta et al., 2019). Feeding procedure excited DCN neurons, while activating DCN neurons suppressed food intake (Low et al., 2021), indicating that DCN contributes to mediate general reward such as food. However, silencing Med^CaMKII^ or activating lobule VI PC^TH+^–Med pathway failed to affect sucrose-induced reinforcements and consumption in mice, suggesting that Med^CaMKII^ and lobule VI PC^TH+^–Med pathway may be likely to specially involved in pathological reward (e.g. METH-related reward in the present study) instead of natural reward.

The DCN expresses dopamine receptors (T. M. Locke et al., 2018), but ISH database of Alan’s mouse brain shows that lobule VI PCs lack Dopa decarboxylase (DDC) and vesicle monoamine transporter Type 2 (VMAT2). Moreover, PC^TH+^ did not exhibit phospho-TH immunostaining in adjacent sections of the cerebellum (Lee et al., 2006). These findings suggest that the TH in PCs may be enzymatically inactive or PCs has no function to synthetize and release dopamine. We thought that lobule VI PC^TH+^ are not dopaminergic neurons, but may be TH-positive interneurons similarly to that in the striatum (Xenias, Ibáez-Sandoval, Koós, & Tepper, 2015). METH has been reported to increase the expression of TH gene and proteins in the cerebellum (Ferrucci et al., 2007). Knocking out TH in PCs impaired behavioral flexibility, social recognition memory, and associative fear learning, but had no deficits in motor, sensory, instrumental learning, or sensorimotor gating functions (T. Locke et al., 2020). Consistently, we found that activating lobule VI PC^TH+^–Med pathway had no effect on motor coordination, while silencing total Med produced motor coordination deficits, indicating that lobule VI PC^TH+^–Med pathway is selectively modulated METH-related reward but not key elements for motor coordination. As such, lobule VI PC^TH+^–Med pathway might be used as a better intervention target for blocking the acquisition (formation) of METH addiction.

In conclusion, our findings revealed a novel cerebellar lobule VI PC^TH+^– Med^CaMKII^ pathway and its potential role in METH-preferred behaviors. Both Med^CaMKII^ and lobule VI PC^TH+^–Med pathway are involved in coding METH-induced CPP behaviors, especially the acquisition (formation) of METH CPP. Importantly, it seemed that suppressing the subpopulations cells of Med innervated by lobule VI PC^TH+^ is better than inactivating total Med to alleviated the METH-preferred behaviors, as no side-effects of activating lobule VI PC^TH+^–Med pathway on motor coordination in mice.

## Materials and Methods

### Animals

C57BL/6 male mice (∼25 g, 8–10 weeks of age) were maintained on a 12-h reverse light-dark cycle with food and water available ad libitum. All mice were handled for three consecutive days prior to experiment. All procedures were carried out in accordance with the National Institutes of Health Guide for the Care and Use of Laboratory Animals and approved by the Institutional Animal Care and Use Committee (IACUC) at Nanjing University of Chinese Medicine.

### Conditioned place preference (CPP)

METH CPP procedures were performed in the TopScan3D CPP apparatus (CleverSys, VA, USA), which is constructed of two distinct chambers separated by a removable guillotine door. The CPP procedure (Pandy et al., 2018) consisted of three phases: the pre-conditioning test (Pre-test), conditioning (CPP training) and post-conditioning test (CPP-test). Mice were allowed a 15-min free exploration to the two chambers as the baseline preference, then equally divided into two sub-groups. Conditioning in METH group was confined to preferred chamber for 45 min paired with a saline (0.2 mL, i.p.) injection, and non-preferred chamber for 45 min paired with a METH (1 mg/kg, i.p.) injection on alternate days (METH on odd days and saline on even days). The saline group was received saline (0.2 mL, i.p.) injection on each training day. During the CPP-test, mice were allowed to freely access the two chambers without any drug treatment for 15 min.

The CPP score was calculated by subtracting the duration spent in saline-paired chamber from the METH-paired chamber. The ⊿CPP score was the test CPP score minus pre-test CPP score.

### Sucrose reinforcement

Sucrose reinforcement experiments were conducted in Skinner box (Harvard Apparatus, USA). During all daily 0.5-h training sessions, each active lever press resulted in the delivery of 0.1 mL 10% sucrose under a FR1 reinforcement schedule with a 5-s timeout. Inactive lever press had no programmed consequences. During the training phase, mice were water restricted but allowed to drink freely for 1h after the daily training.

### Rota-rod task

Motor coordination and balance were measured by the accelerating rota-rod test (4 rpm to 40 rpm in 5min) (Chen et al., 2018), in which mice were placed on the 3-cm diameter rota-rod cylinder (Ugo Basil, Italy) during the 5-min test. Mice were initially trained to maintain themselves in a neutral position on the rod, and the latency to fall off the rota-rod was calculated.

### Locomotor activity test

General locomotor activity was measured in an open field arena. Mice were placed in the chamber and allowed to freely explore for 10 min. The total distance traveled was recorded and analyzed by an automated detection system (TopScan, USA).

### Immunofluorescence

The mice were perfused with 0.9% saline followed by 4% paraformaldehyde (PFA) in PBS buffer. The brains were removed and post-fixed in 4% PFA at 4 °C overnight, then transferred to 30% (w/v) sucrose. Frozen sections (30 μm) were cut on a cryostat (Leica, Germany). The sections were incubated with the primary antibodies overnight at 4°C, followed by the corresponding fluorophore-conjugated secondary antibodies for 1.5 h at room temperature. The following primary antibodies were used: mouse anti-NeuN (1:1000, 94403S, Cell Signaling Technology, RRID: AB_2904530), rabbit polyclonal anti-NeuN (1:500, 2407S, Cell Signaling Technology, RRID: AB_2651140), mouse anti-CaMKII (1:150, sc-13141, Santa Cruz, RRID:AB_626789), rabbit polyclonal anti-TH (1:250, BM4568, Boster), mouse anti-Calbindin (1:800, ab75524, abcam, RRID: AB_1310017), rabbit polyclonal anti-c-Fos (1:500,226003, Synaptic Systems, RRID: AB_2231974). The secondary antibodies were used: Alexa Fluor 555-labeled donkey anti-mouse IgG (1:500, A322773, Invitrogen, RRID: AB_2762848), Alexa Fluor 555-labeled donkey anti-rabbit (1:500, A32794, Invitrogen, RRID: AB_2762834), Alexa Fluor 488-labeled donkey anti-mouse (1:500, A21202, Invitrogen, RRID: AB_141607), Alexa Fluor 488-labeled donkey anti-rabbit (1:500, A32790, Invitrogen, RRID: AB_2762833). The images were captured by Leica DM6B upright digital research microscope (Leica, Germany) and THUNDER imaging systems TCS SP8 (Leica, Germany).

### Stereotaxic surgery

All viruses were generated and packaged by BrainVTA (Wuhan, China). Mice were head-fixed with a stereotactic frame (RWD, Shenzhen, China) under isoflurane anesthesia (2% induction, 0.5% maintenance). 100 nL of viruses or Fluorogold (FG) were injected using a glass micropipette attached to an infusion pump (Drummond, Alabama, USA) over 5 min at a rate of 20 nL/min. We waited for 10 min both before and after each injection. The coordinates used here were: Med (AP, −6.40 mm; ML, ±0.75 mm; DV, −3.55 mm), lobule VI (AP, −6.80 mm; ML, 0 mm; DV, −1.20 mm). For retrograde tracing, 100 nL FG (Fluorochrome, 4%) was injected unilaterally in the Med. For anterograde tracing, 100 nL of a 1:1 volume mixture of AAV2/9-mTH-cre and AAV2/9-L7-DIO-mCherry was injected unilaterally in the lobule VI. For the tettox experiments, 100 nL of a 1:1 volume mixture of *AAV2/9-CaMKII-cre* and *AAV2/9-EF1α-DIO-tettoxlc-mCherry* (or *AAV2/9-EF1α-DIO-EGFP*) was injected bilaterally in the Med. For local infusion of CNO, pedestal guide cannulas (27-gauge, RWD Life Science) were implanted bilaterally 1 mm above the Med, along with 100 nL of a 1:1 volume mixture of *AAV2/9-mTH-cre and AAV2/9-L7-DIO-hM3Dq-mCherry*.

### Patch clamp

Slices preparation were performed as previously described (Ge et al., 2021). Mice were deeply anesthetized with isoflurane (RWD, China) and perfused with the ice-cold cutting solution. Sagittal slices containing the lobule VI were cut at 200 μm thickness using a vibratome in 4°C cutting solution. The slices were transferred to 37 °C cutting solution for 9 min, then transferred to holding solution to allow for recovery at room temperature for 1 h before recordings. During electrophysiological recordings, the brain slice was continuously perfused with oxygenated artificial CSF (aCSF) maintained at 30°C by an in-line solution heater (TC-324C, Warner Instruments, USA). Loose-patch electrodes were filled with aCSF, and access resistance was maintained at 20–50 MΩ throughout the experiment. Recordings were performed under current-clamp mode with 0 holding current. All signals were filtered at 4 KHz, amplified at 5x using a MultiClamp 700B amplifier (Molecular Devices, USA), digitized at 10 kHz with a Digidata 1440A analog-to-digital converter (Molecular Devices). All data were analyzed with Clampfit 10.6 software (Molecular Devices, USA).

### Statistical analysis

Statistical analysis was carried out using GraphPad Prism 9.0 software. The data are presented as the mean ± s.e.m. Statistical significance was set as \**P* < 0.05, \*\**P* < 0.01 and \*\*\**P* < 0.001, as determined by two-tailed unpaired t-test, one-way repeated measure (RM) ANOVAs, two-way repeated-measures analysis of variance (ANOVAs) followed by Bonferroni’s post-hoc test.

## Acknowledgments

This work is supported by National Natural Science Foundation of China (No. 82071495), Natural Science Foundation of Jiangsu Province, China (No. BK20201398), and the Open Project of Chinese Materia Medica First-Class Discipline of Nanjing University of Chinese Medicine (2020YLXK004).

## Author contributions

Ge F, Wang Z, Wang G, Yang P, Wen J, Pan W and Yu W were responsible for the experiments. Ge F and Chang J were responsible for formal analysis. Cai Q and Fan Y assist the data analysis. Guan X and Ge F wrote the manuscript. Guan X developed the overall concept.

## Supplement Figure Legends

**Figure supplement 1.**
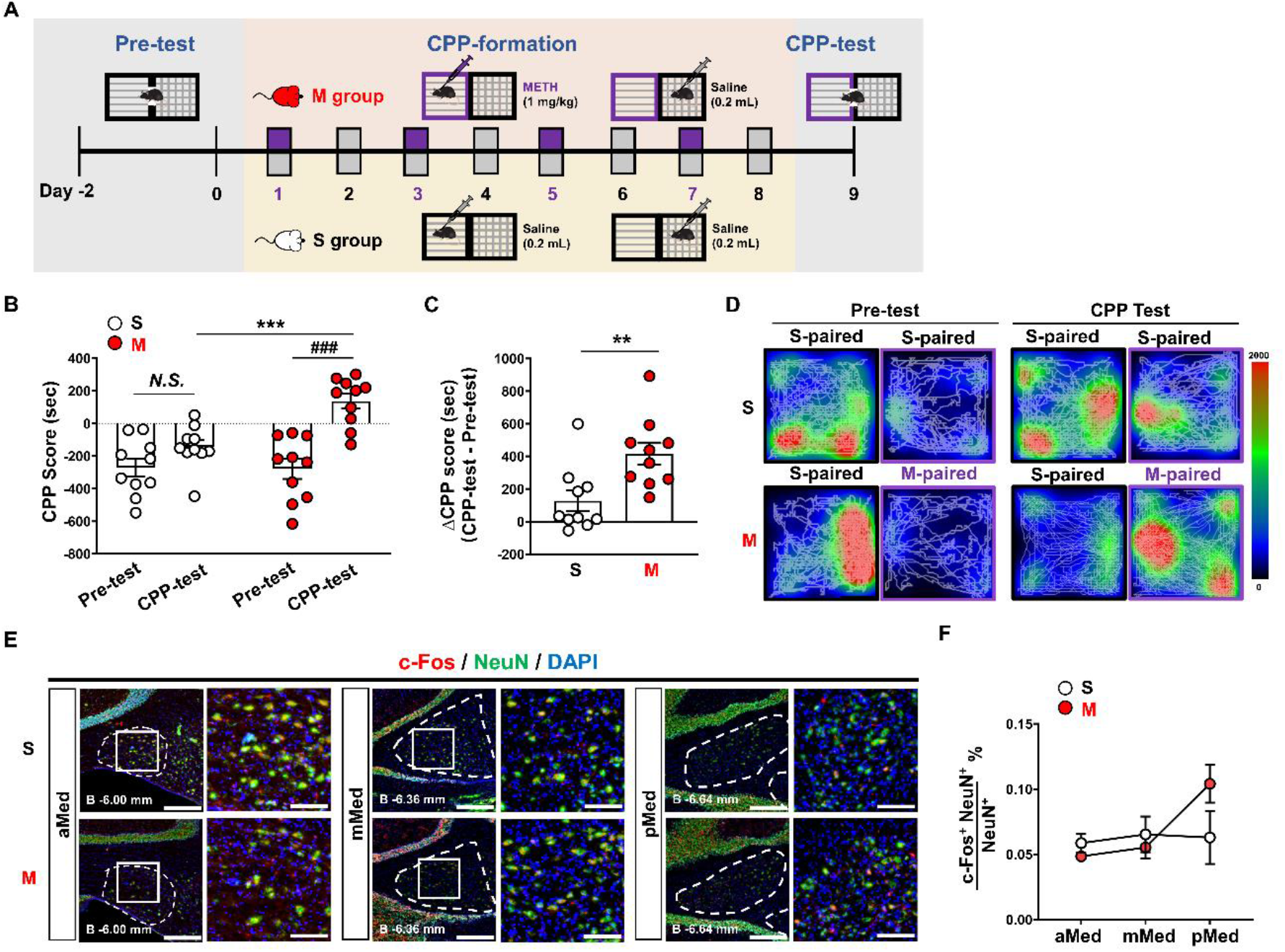
METH CPP test has no effect on the activities of Med. (**A**) Scheme of METH CPP test model. (**B**) CPP scores during the pre-test and CPP test. Two-way RM ANOVA. F _treatment × session_ (1,18) =9.689, *P* = 0.0060. ^###^*P* < 0.001 versus Pre-test of METH group. *^**^*P* < 0.001 versus saline group during the CPP-test. n = 10 per group. (**C**) ΔCPP score (CPP test minus pre-test) of the two group. Two-tailed unpaired t test, t = 3.113, df = 18, ^**^*P* = 0.0060 versus saline group. n = 10 per group. (**D**) Single trial example during the pre-test and CPP test. (**E**) Immunohistochemistry for c-Fos/NeuN/DAPI in the Med subregions following METH CPP test. Scale bar, 300 μm (left) and100 μm (right). (**F**) Percentage of c-Fos and NeuN double positive neurons in the Med following METH CPP formation. Two-way RM ANOVA. F _treatment × session_ (2,20) = 0.8325, *P* = 0.4495. ^*^*P* < 0.05 versus saline group. n = 6 per group.

**Figure supplement 2.**
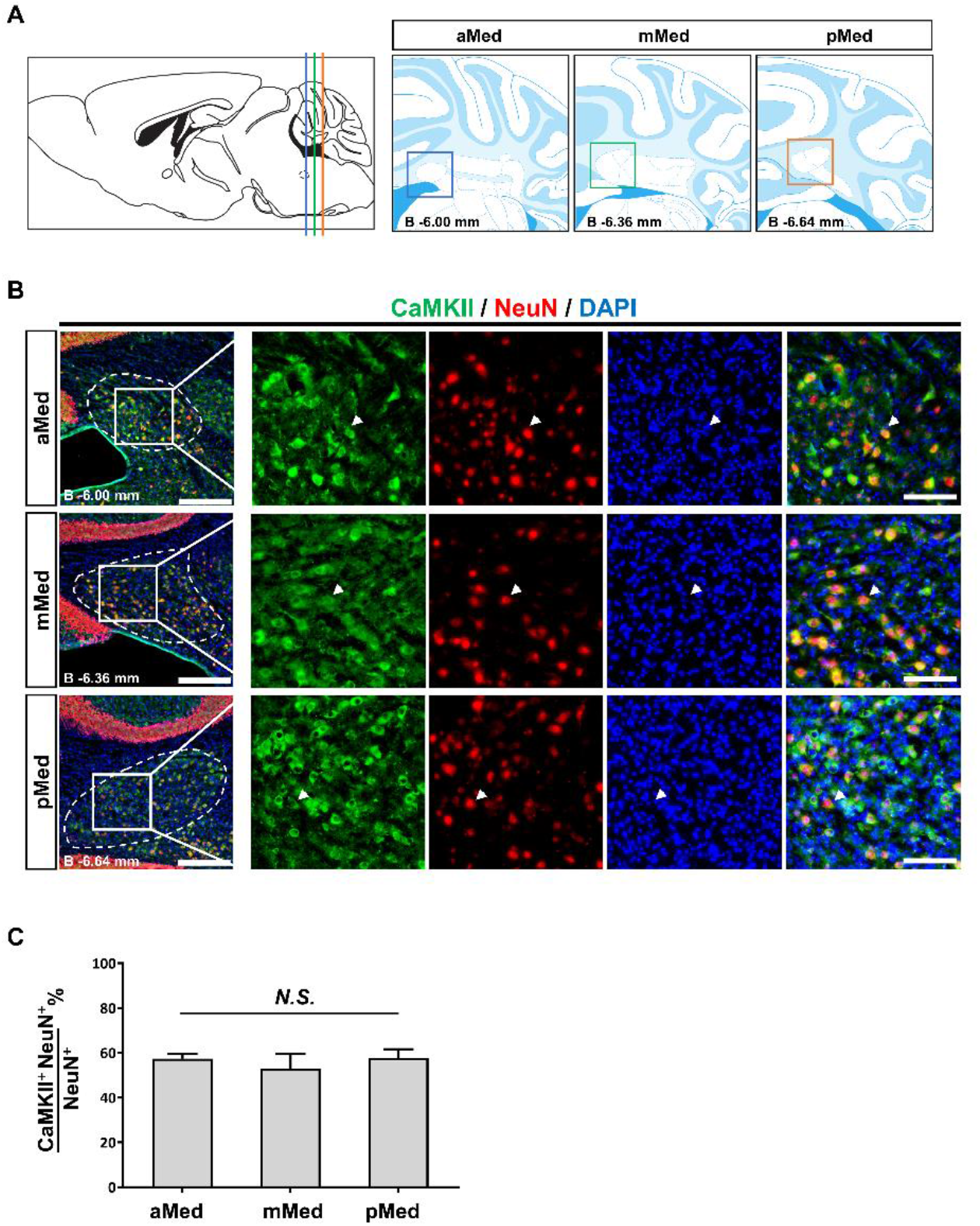
Med neurons are predominantly CaMKII-positive glutamatergic neurons. (**A**) Schematics of Med subregions. (**B**) Immunohistochemistry for CaMKII/NeuN/DAPI in the Med subregions of naïve mouse. Scale bar, 300 μm (left) and 100 μm (right). (**C**) Percentage of CaMKII and NeuN double positive cells in the Med subregions. One-way ANOVA. F (2,3) = 0.3197, *P* = 0.7484. n = 2.

**Figure supplement 3.**
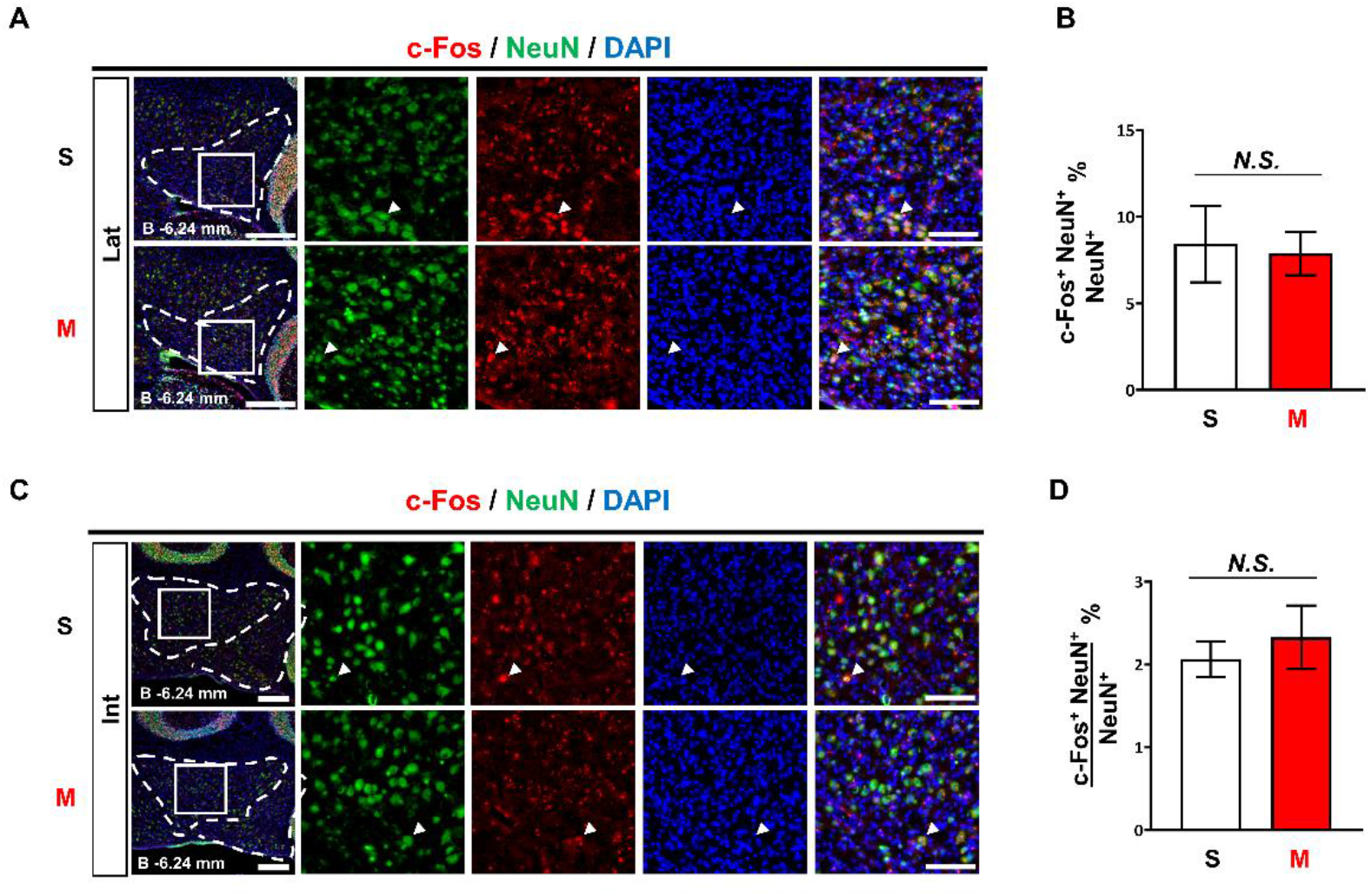
METH CPP formation had no effect on the activities of cerebellar Lat and Int. (**A**) Immunohistochemistry for c-Fos/NeuN/DAPI in the Lat following METH CPP formation. Scale bar, 300 μm (left) and 100 μm (right). (**B**) Percentage of c-Fos and NeuN double positive cells in the Lat following METH CPP formation. Two-tailed unpaired t test, t = 0.2148, df = 10, *P* = 0.8343 versus saline group. n = 6 per group. (**C**) Immunohistochemistry for c-Fos/NeuN/DAPI in the Int following METH CPP formation. Scale bar, 300 μm (left) and 100 μm (right). (**D**) Percentage of c-Fos and NeuN double positive cells in the Int following METH CPP formation. Two-tailed unpaired t test, t = 0.6011, df = 10, *P* = 0.5611 versus saline group. n = 6 per group.

**Figure supplement 4.**
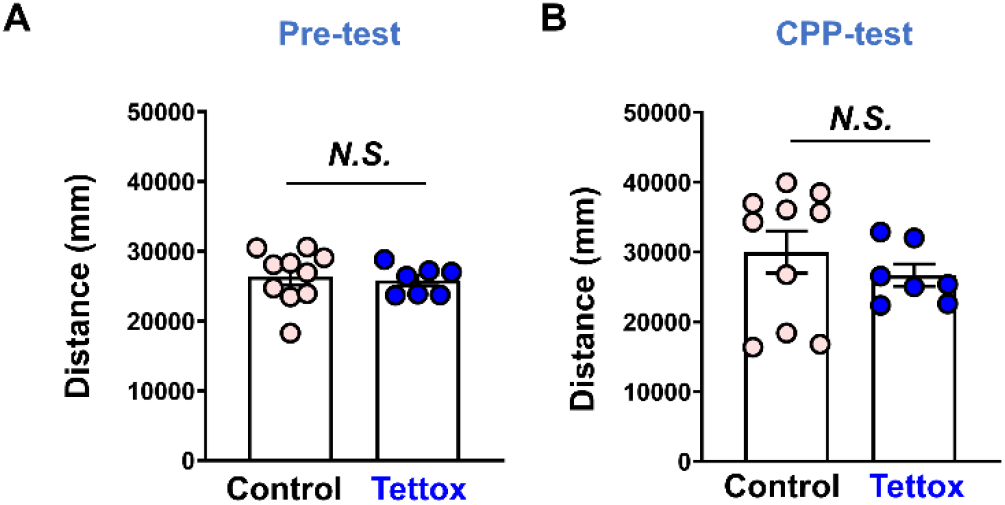
Sustained silencing Med glutamatergic neurons has no effect on distance travelled during the pre-test and CPP test. (**A**) Distance travelled during the pre-test. Two-tailed unpaired t test, t = 0.3521, df = 15, ^**^*P* = 0.7297 versus control group. Control group, n = 10; Tettox group, n = 7. (**B**) Distance travelled during the CPP test. Two-tailed unpaired t test, t = 0.8539, df = 15, *P* = 0.4066 versus control group. Control group, n = 10; Tettox group, n = 7.

## Graphical Abstract

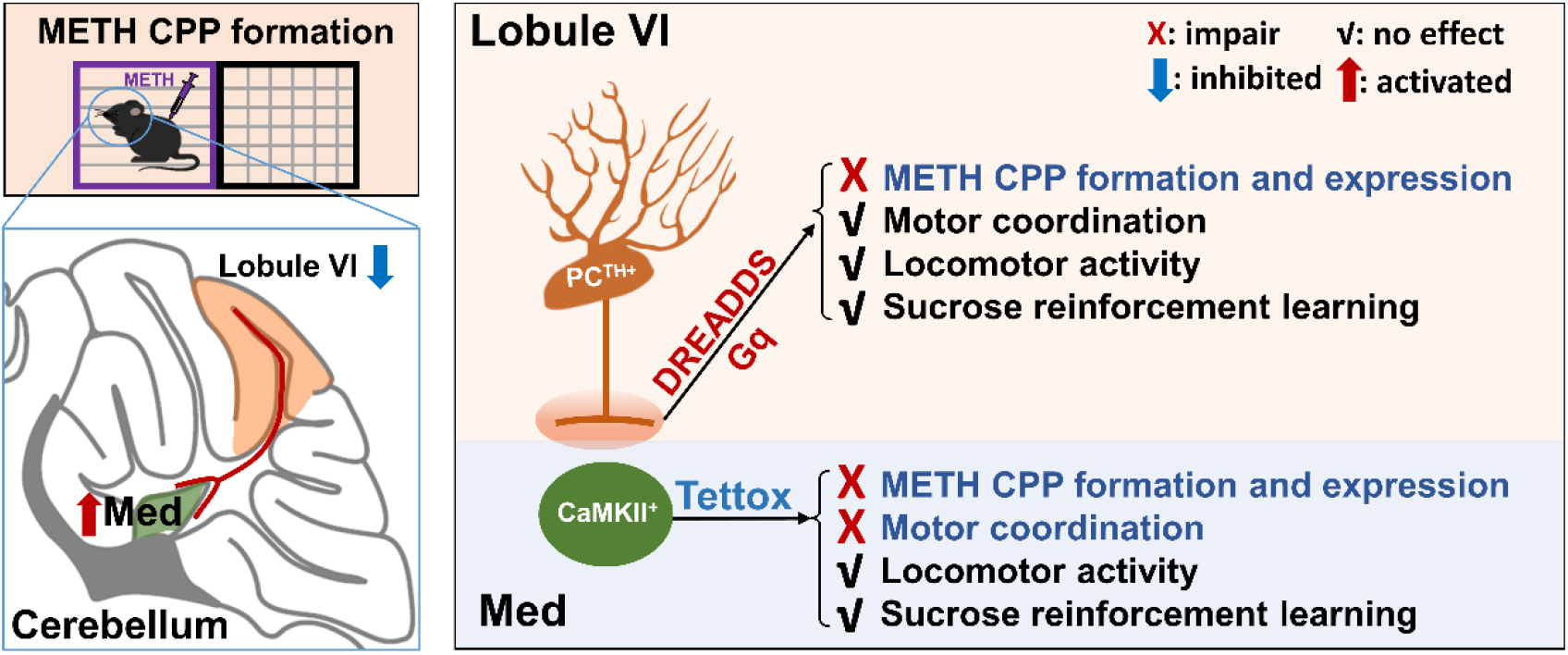

